# Identity signaling, identity reception and the evolution of social recognition in a Neotropical frog

**DOI:** 10.1101/2021.04.25.441331

**Authors:** James P. Tumulty, Zachary K. Lange, Mark A. Bee

## Abstract

Animals recognize familiar individuals to perform a variety of important social behaviors. Social recognition is often mediated by communication between signalers who produce signals that contain identity information and receivers who categorize these signals based on previous experience. We tested two hypotheses about adaptations in signalers and receivers that enable the evolution of social recognition using two species of closely related territorial poison frogs. Male golden rocket frogs (*Anomaloglossus beebei*) recognize the advertisement calls of conspecific territory neighbors and display a “dear enemy effect” by responding less aggressively to neighbors than strangers, while male Kai rocket frogs (*A. kaiei*) do not. Our results did not support the identity signaling hypothesis: both species produced advertisement calls that contain similar amounts of identity information. Our results did support the identity reception hypothesis: both species exhibited habituation of aggression to playbacks simulating the arrival of a new neighbor, but only golden rocket frogs showed renewed aggression when they subsequently heard calls from a different male. These results suggest that an ancestral mechanism of plasticity in aggression common among frogs has been modified through natural selection to be specific to calls of individual males in golden rocket frogs, enabling a social recognition system.

## 1. Introduction

Humans and a diversity of other animals exhibit social recognition [1–3], the act of recognizing other individuals based on previous experiences and assigning them to one or more social categories (e.g., “neighbor”, “offspring”, or “nest-mate”). Across animals, recognized social categories vary along a continuum of specificity, ranging from species recognition at the most general end of the continuum to the recognition of particular individuals at the most specific [4]. Social recognition often involves learning the features of another individual’s communication signals, such as their visual markings (e.g., [5]) or voices (e.g., [6]). Two non-mutually exclusive hypotheses explain how signalers and receivers in a communication system can be targets of selection for social recognition. The *identity signaling hypothesis* states that, if there is a net benefit of being recognized, selection acts on signalers and favors an increase in the individual distinctiveness of signals, such that greater information about a signaler’s identity facilitates correct recognition [7,8]. This hypothesis predicts more identity information in the signals of species with social recognition than in those without. Provided identity information exists, receivers must be able to perceive individual differences in signals, remember the signal properties of familiar individuals, and employ decision rules for discriminating between individuals that belong to different social categories [2,9,10]. According to what we refer to here as the *identity reception hypothesis*, social recognition evolves in response to selection on pre-existing mechanisms of learning and discrimination in receivers to increase the specificity of learned social categories. This hypothesis predicts that individuals of species that exhibit social recognition learn to discriminate narrower social categories of conspecifics than those of species that do not exhibit social recognition. Comparative studies of closely related species that differ in social recognition are key to testing the predictions of both hypotheses [11–13], but few such studies exist, in part, because closely related species (e.g., songbirds [14,15]) often share similar social recognition abilities.

Here, we tested the identity signaling and identity reception hypotheses in a field study of two closely related Neotropical poison frogs (Aromobatidae, Dendrobatoidea), the golden rocket frog (*Anomaloglossus beebei*) and the Kai rocket frog (*Anomaloglossus kaiei*). These two territorial species differ in a form of social recognition in which territory holders display lower levels of aggression toward familiar neighbors compared with strangers, a behavior known as the “dear enemy effect” [14,16,17]. In many animals [14], including songbirds [15] and a small number of frog species [18], neighbor recognition and the dear enemy effect are based on a receiver’s ability to learn individually distinctive features of their neighbor’s vocalizations. Golden rocket frogs recognize and respond less aggressively to the individually distinctive advertisement calls of neighbors compared with those of strangers [17,19,20]. In contrast, Kai rocket frogs are similar to other poison frogs studied to date [21,22] in being equally aggressive to neighbors’ and strangers’ calls [17]. Neighbor recognition and the dear enemy effect likely evolved in golden rocket frogs in response to ecological and social conditions associated with breeding in spatially clumped bromeliads (“phytotelm breeding”), which puts males in closer proximity for longer periods of time than does the terrestrial reproductive ecology of Kai rocket frogs [17]. Kai rocket frogs defend territories on the forest floor and transport tadpoles from terrestrial egg clutches to pools of water [17]. Terrestrial breeding is ancestral in the genus *Anomaloglossus* (Fig. 1a), suggesting that phytotelm breeding and any associated behaviors are evolutionarily derived in golden rocket frogs. Therefore, by comparing the communication systems of Kai rocket frogs (representing the ancestral state) and golden rocket frogs (representing the derived state) we can investigate evolutionary changes in signalers and receivers associated with the evolution of a social recognition system.

**Figure 1.**
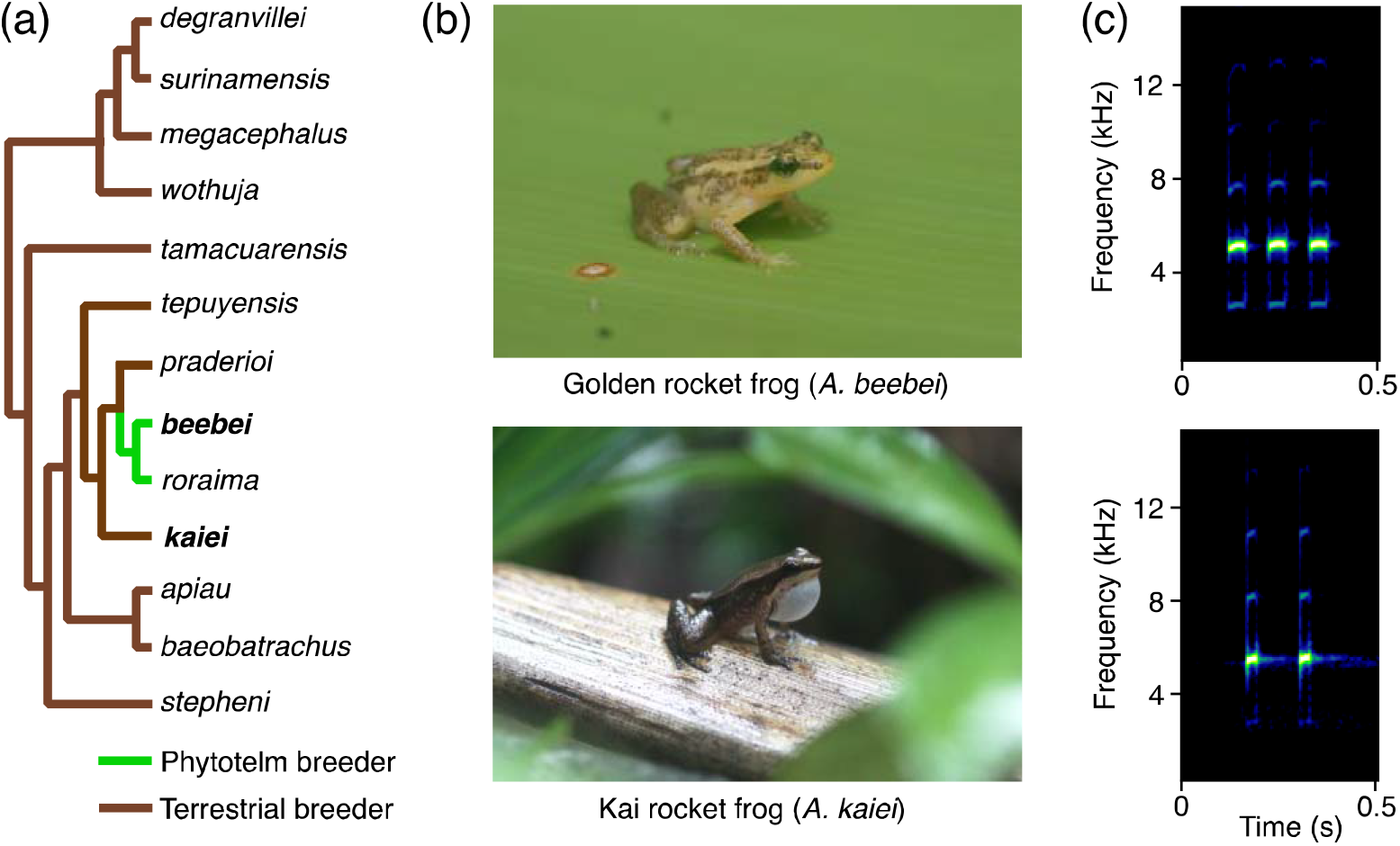
Two closely related species were the subjects of this comparative study. (a) Cladogram of the genus *Anomaloglossus*, redrawn and trimmed from [32], with branch colors indicating reproductive ecology (phytotelm or terrestrial breeder) demonstrating that phytotelm breeding is evolutionarily derived in golden rocket frogs (*A. beebei*). On this cladogram, terrestrial breeders consist of species that breed on the forest floor and those that breed along streams. Note: branch lengths are not meaningful. (b) Photographs of a male golden rocket frog and a male Kai rocket frog. (c) Spectrograms of the advertisement calls of golden rocket frogs (top) and Kai rocket frogs (bottom).

According to the identity signaling hypothesis, we predicted greater identity information in the calls of golden rocket frogs compared with Kai rocket frogs. We tested this prediction by measuring acoustic properties of advertisement calls and statistically comparing information content and patterns of individual variation in these properties across species. According to the identity reception hypothesis, we also predicted golden rocket frogs would learn to recognize narrower, individual-specific social categories compared with Kai rocket frogs. As a well-known form of learning, stimulus-specific habituation has been identified as a potential mechanism underlying the reduced aggression typical of the dear enemy effect [23–26]. Previous work in fish [23], songbirds [24], and frogs [18] has demonstrated that territorial males become less aggressive (i.e., habituate) upon repeated exposures to real or simulated unfamiliar individuals, but that renewed aggressive responses can be elicited by changing relevant features of the stimulus, such as its vocal identity. Therefore, we performed a field playback experiment based on the habituation-discrimination paradigm to compare between the two species the specificity of social categories that are learned through repeated exposure to a new neighbor.

## 2. Materials and methods

### (a) Study site and species

We studied golden and Kai rocket frogs in Kaieteur National Park, Guyana, from 2013 to 2017. We monitored males by regularly surveying sites where members of each species were locally abundant, and identified individual males through a combination of unique toe clips and photographs of dorsal color patterns (see [17] for more details). A maximum of one toe per foot was clipped using a pair of ethanol and flame-sterilized spring scissors, following established protocols [27]. Golden rocket frogs call from and defend territories in giant tank bromeliads, while Kai rocket frogs call from and defend territories on the forest floor [17,19,28–30]. Advertisement calls typically consist of a series of 3 pulses (range 1-6) for golden rocket frogs or 2 pulses (range 1-2) for Kai rocket frogs (Fig. 1c, Fig. S1 [19,20,28,31]). Both species have distinct aggressive calls consisting of longer trains of pulses, as well as shortened “pseudo-aggressive” calls (Fig. S1 [17]). Males of both species respond to conspecific intruders with aggressive calls followed by rapid approach movements, wrestling, and chasing. Components of this territorial response, including aggressive calls and approach movements, can be elicited in response to playbacks of conspecific advertisement calls.

### (b) Identity signaling hypothesis

#### (i) Acoustic recordings and measurements

We analyzed the patterns of individual variation in a large set of recorded advertisement calls of previously identified territorial males of each species. Recordings were made from a distance of 0.5-1 m using a directional microphone (Sennheiser ME-66) and digital recorder (Marantz PMD-620; 44.1 kHz sampling rate, 16-bit resolution). After recording, we caught the frog to confirm its identity and measured the air temperature at the site of calling (ranges: 22.8 to 25.8°C for golden rocket frogs and 21.2 to 25.2°C for Kai rocket frogs). We measured the temporal and spectral properties (see Fig. S2) of 20 advertisement calls from each of 20 males per species (800 calls total) using Raven Pro 1.5 (Cornell Lab of Ornithology). We measured temporal properties of calls (call duration, call interval, and call period), temporal properties for each pulse within a call (pulse duration, pulse interval, and pulse period), temporal properties describing pulse shape (pulse rise time, pulse 50% rise time, pulse fall time, and pulse 50% fall time), and dominant frequency. We limited analyses of golden rocket frog calls to those with four or fewer because males rarely produced calls with more pulses (2% of calls; see also [30,31]).

#### (ii) Statistical analysis

Prior to the following statistical analyses, we removed any temperature-dependent variation in acoustic properties by using linear regression to standardize values of all call properties to the mean temperatures at which recordings were made (23.5°C for golden rocket frogs and 22.7°C for Kai rocket frogs). Such standardization removes variation due to a known environmental influence on frog calls that is not due to reliable individual (phenotypic) differences [33–35].

For each species and acoustic property, we calculated a mean and standard deviation for each individual, the grand mean and standard deviation based on the average values for 20 individuals, and the “potential for individual coding” (PIC, [36]), which was the ratio of the among-individual coefficient of variation (CV_a_ = grand SD/grand mean × 100%) to the within-individual coefficient of variation (CV_w_ = individual SD/individual mean × 100%). The proportion of variance explained by individual differences was calculated as the effect size (partial η^2^) of “individual” as a random effect in a separate ANOVA conducted for each property.

For multivariate measures of identity information, we incorporated all call properties but restricted our analysis to calls with at least three pulses for golden rocket frogs (*n* = 350 of 400 calls) and at least two pulses for Kai rocket frogs (*n* = 383 of 400 calls), because these methods require complete data sets. The data were centered, scaled, and subjected to a principal components analysis to yield uncorrelated transformed variables using the ‘prcomp’ function in R (R Foundation for Statistical Computing). We retained all components that cumulatively explained greater than 99% of the variation in the data set (17 for golden rocket frogs and 12 for Kai rocket frogs). Note, conclusions did not change when running subsequent analyses using more restrictive sets of principal components (see Table S3). Principal components were used to compute Beecher’s information statistic (*H_s_*), which measures identity information as a signal’s ability to reduce uncertainty about the identity of the signaler [7,36]. We also included these principal components as input variables in a linear discriminant analysis using the ‘lda’ function in the ‘MASS’ package in R and used a leave-one-out cross-validation procedure to evaluate the accuracy with which calls could be assigned to the correct individual. To test for a species difference in discrimination accuracy, we modelled whether or not a call was assigned to the correct individual using a logistic mixed-effects model that included species as a main effect and individual as a random effect. The statistical significance of species differences was tested using likelihood ratio tests. Because territorial males of both species typically had only a small number of adjacent neighbors [17], we also ran the linear discriminant analysis on 100 randomly selected groups of five individuals to examine the statistical discrimination of individuals using a more ecologically relevant discrimination task [33,34].

### (c) Identity reception hypothesis

#### (i) Experimental design

We conducted a habituation-discrimination playback experiment in the field using 20 territorial males of each species as subjects (total *n* = 40). Testing protocols for the two species were designed to be as similar as possible while accommodating the constraints of natural species differences in call rate and intermale distances. Each test with a subject consisted of a habituation phase followed immediately by a discrimination phase. During the habituation phase, we repeatedly presented subjects with a 1-min habituation stimulus consisting of a set of calls produced by a conspecific male. During a 4-min stimulus period, the 1-min habituation stimulus was presented twice consecutively (i.e., 2 min of simulated calling) followed by 2 min of silence. Provided the subject responded aggressively during at least one of the first three stimulus periods, we continued repeating the 4-min stimulus period during the habituation phase until the subject met pre-established habituation criteria, which were no aggressive calls and no approach movements towards the speaker for three consecutive stimulus periods. Upon meeting these habituation criteria, subjects were randomly assigned to either a ‘different frog’ experimental treatment (*n* = 10 per species) or a ‘same frog’ control treatment (*n* = 10 per species) during the subsequent discrimination phase, which lasted for three additional stimulus periods. In both treatments, subjects heard a novel set of calls they had not heard previously. In the different frog treatment, the novel stimulus consisted of a new set of calls recorded from a different male than the one who produced the calls used as the habituation stimulus. In the same frog treatment, the new set of calls was produced by the same male who produced the calls used for the habituation stimulus. Hence, only subjects in the different frog treatment experienced a change in the identity of the simulated male. Stimuli were fully replicated such that each stimulus came from a unique male, except for the two paired stimuli from the same male for the same frog treatment.

#### (ii) Experimental protocol

We used Adobe Audition 1.5 (Adobe Systems Inc.) to create acoustic stimuli (WAV files, 44.1 kHz sampling rate, 16-bit resolution) from our field recordings. We standardized the call rate of each stimulus to match the mean rate at which males of each species naturally call (1 call/2.5 s for golden rocket frogs and 1 call/s for Kai rocket frogs) and created 1-minute stimuli consisting of 23 unique calls for golden rocket frogs or 59 unique calls for Kai rocket frogs by inserting an appropriate amount of silence between consecutive calls. Thus, for each species, our stimuli incorporated the natural variation in calls that would occur over an average 1-minute period of calling. Stimuli were high-pass filtered above 2 kHz to minimize background noise and normalized to 80% relative amplitude. File names of stimuli were changed and arranged in random order on the playback device so that experimenters were blind to treatment when conducting playback tests and scoring responses.

Prior to a playback test, we located calling males and placed a speaker (Saul Mineroff Electronics) facing the male from a direction without close neighbors at a distance of 1.5 m for golden rocket frogs and 2-3 m for Kai rocket frogs. The difference in speaker distance was required to incorporate natural species differences in inter-male distance into our experimental design [17]. Stimuli were broadcast using an iPod (Apple Inc.). We calibrated the speaker to produce stimuli at 80 dB sound pressure level at 1 m (SPL re 20 μPa, fast RMS, A-weighted) in suitable habitat for each species using a Type 2 digital sound level meter (407764, Extech). This SPL is within the range of natural variation for both species’ calls. The frequency response of the playback system was flat (± 2.4 dB) across the dominant frequency range (4.6-5.8 kHz) of both species’ calls.

During each minute of the test, we recorded four measures of aggression: 1) number of aggressive calls, 2) number of pseudo-aggressive calls, 3) approach distance, and 4) the subject’s closest position to the speaker. We recorded the acoustic responses of males during a test using the same recording equipment described above, and call counts were later confirmed using these recordings. We used local landmarks (e.g., leaves, logs) to note the subject’s location throughout the test and, afterwards, measured the distances from the speaker to each noted location to provide approach distance and closest position to the speaker measurements. We also measured the distance from the subject’s original location to its nearest neighbor. If the neighbor was not calling at the time, we measured the distance to the approximate center of the neighbor’s territory. We measured the SPL of both stimuli at the male’s original location; all habituation and discrimination stimuli differed by less than 2.5 dB. We aborted the test if the subject did not show any aggressive behaviors during the first three stimulus periods of the habituation phase, started courting a female, or interacted aggressively with a real male.

#### (iii) Statistical analysis

For each subject, we summarized the data in blocks consisting of three stimulus periods. Within each block, we assessed a subject’s aggressiveness by totaling the numbers of aggressive and pseudo-aggressive calls, summing the approach distances, and determining the subject’s closest location to the speaker. To control for slight variation in speaker placement, we expressed each subject’s closest location to the speaker as a proportion of the speaker distance at the start of the test. Because subjects took different amounts of time to habituate, we analyzed just the first and last block of the habituation phase, as well as the single block of the discrimination phase. Separately for each species, we computed an aggression index as the first principal component from a principal components analysis on all four measures of aggression (centered and scaled) using the ‘prcomp’ function in R (factor loadings in Table S8). To compare treatment differences in recovery of aggression during the discrimination phase, we modeled the aggression index of males in the last block of the habituation phase versus the discrimination phase using linear mixed effects models implemented through the ‘lmer’ function in the ‘lme4’ package. This model included block (last block of habituation phase vs. single block of discrimination phase) and treatment (different frog vs. same frog) as fixed effects and individual as a random effect. To evaluate whether any recovery of aggression during the discrimination phase was dependent on the identity of the simulated intruder, we tested for a significant interaction between block and treatment. We then ran separate tests within treatment to test for increases in aggression from the last block of the habituation phase to the discrimination phase again using linear mixed effects models with individual as a random effect. Significance of fixed effects was evaluated using likelihood ratio tests. We used Wilcoxon signed-rank tests to compare the decrease in aggression during the habituation phase and Wilcoxon rank sum tests to compare habituation time between species because both data sets did not meet parametric assumptions. We also used Spearman’s rank-order correlations to evaluate the relationship between habituation time and nearest neighbor distance.

## 3. Results

### (a) Identity signaling hypothesis

Both species’ calls exhibited similar patterns of individual variation, as indicated by the broad similarity in PIC values (Tables 1, S4, S5). In both species, the gross temporal features of pulses within calls (pulse duration, pulse interval, and pulse period) and dominant frequency had the largest PIC values (>2.0) and effect sizes (>0.5) for individual differences (Tables 1, S4, S5). Properties describing temporal features of calls (call duration, number of pulses per call, call interval, and call period) and those describing the fine scale shapes of individual pulses within calls (rise time, fall time, 50% rise time, and 50% fall time) had lower PIC values that were closer to 1.0 as well as smaller effect sizes (Tables 1, S4, S5).

**Table 1.** Descriptive statistics of within- and among-individual variation in acoustic properties of advertisement calls, including estimates of the potential for individual coding (PIC) and effect sizes associated with individual differences (partial *η^2^*).

| Call property | Golden rocket frogs |  |  |  | Kai rocket frogs |  |  |  |
| --- | --- | --- | --- | --- | --- | --- | --- | --- |
| | CV<sub>a</sub> | Mean<br>CV<sub>w</sub> | PIC | Partial $\eta^2$ | CV<sub>a</sub> | Mean<br>CV<sub>w</sub> | PIC | Partial $\eta^2$ |
| Call duration | 19.4 | 16.5 | 1.2 | 0.54 | 10.8 | 11.4 | 0.9 | 0.31 |
| No. pulses | 16.5 | 12.8 | 1.3 | 0.57 | 4.8 | 5.3 | 0.9 | 0.20 |
| Call interval | 32.6 | 39.1 | 0.8 | 0.09 | 43.1 | 40.1 | 1.1 | 0.22 |
| Call period | 29.5 | 36.8 | 0.8 | 0.08 | 36.7 | 34.5 | 1.1 | 0.22 |
| **Pulse duration\*** | **9.3** | **4.7** | **2.1** | **0.81** | **11.5** | **5.7** | **2.0** | **0.79** |
| **Pulse interval\*** | **9.3** | **4.7** | **2.0** | **0.79** | **6.4** | **3.1** | **2.1** | **0.59** |
| **Pulse period\*** | **6.3** | **2.9** | **2.3** | **0.79** | **5.7** | **2.3** | **2.5** | **0.65** |
| Pulse rise time* | 29.8 | 39.4 | 0.8 | 0.35 | 44.8 | 41.0 | 1.1 | 0.54 |
| Pulse fall time* | 27.1 | 32.1 | 0.9 | 0.40 | 24.6 | 27.5 | 0.9 | 0.46 |
| Pulse 50% rise time* | 55.1 | 40.7 | 1.4 | 0.48 | 47.3 | 33.3 | 1.5 | 0.53 |
| Pulse 50% fall time* | 58.6 | 63.0 | 1.0 | 0.23 | 64.2 | 52.6 | 1.2 | 0.56 |
| **Dominant frequency\*** | **4.2** | **1.6** | **2.8** | **0.71** | **3.3** | **1.3** | **2.6** | **0.81** |

Properties with PIC $\geq 2$ are highlighted in bold.
\*To simplify presentation, we present the averages of descriptive statistics of acoustic properties measured on multiple pulses within a call. Descriptive statistics for each pulse presented separately can be found in the supporting information.

Both species’ calls contained similar amounts of identity information (golden rocket frogs, *H_s_* = 5.09 bits; Kai rocket frogs, *H_s_* = 4.57 bits). The slightly higher identity information in the calls of golden rocket frogs was likely due to the greater number of pulses in their calls (Figure 1a). A separate analysis restricted to the first two pulses of golden rocket frog calls yielded a lower estimate of identity information (*Hs* = 4.31 bits; Table S3) that was more similar to the estimate for Kai rocket frogs, which have two-pulse calls. We found no species difference in the success with which linear discriminant analysis assigned calls to the correct individual when we used the full sample of 20 individuals per species (golden rocket frogs: 86%; Kai rocket frogs: 84%; *X^2^* = 0.21, *p* = 0.646). This outcome also held when we restricted these analyses to 100 random samples of 5 individuals per species (golden rocket frogs, mean = 94%, range 85-100%; Kai rocket frogs, mean = 94%, range 80-100%) (Fig. 2).

**Figure 2.**
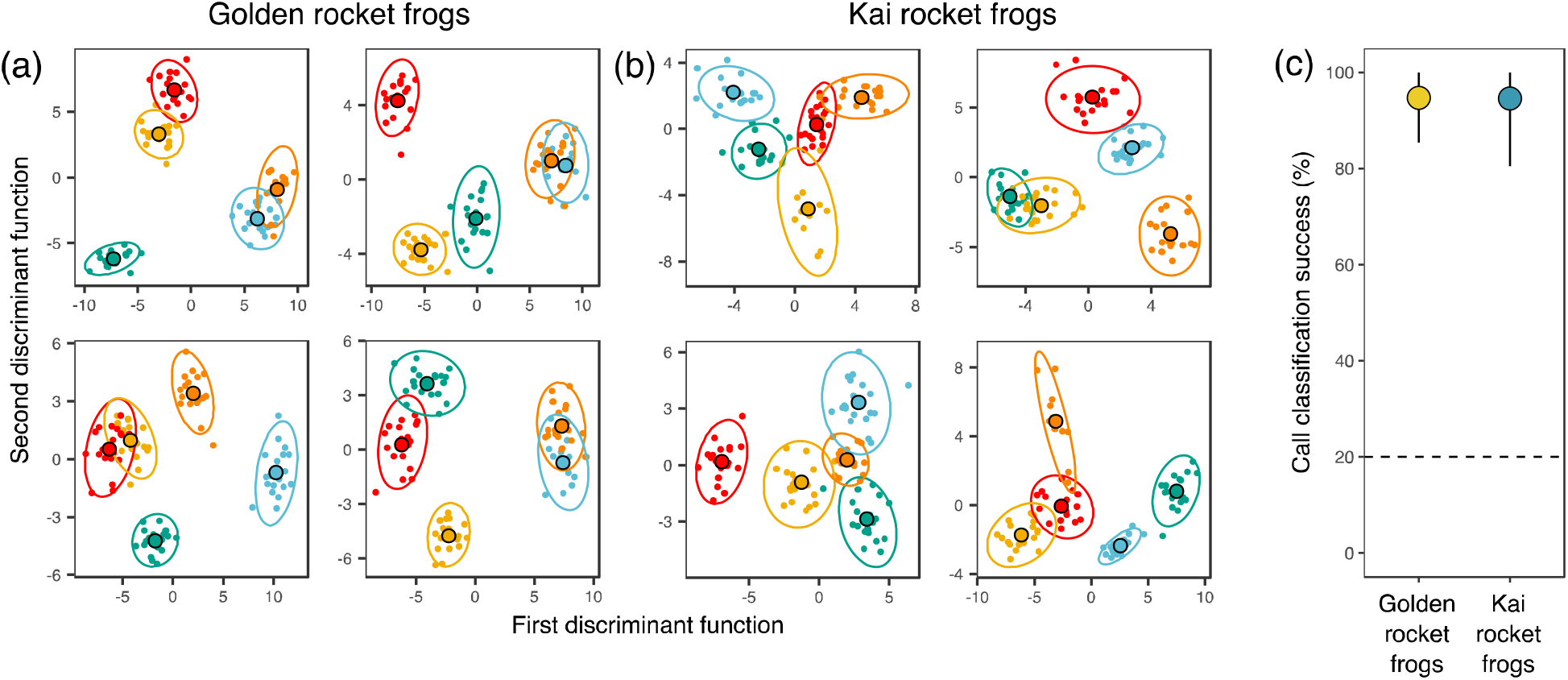
Individual differences in the calls of both species can be statistically discriminated with similar accuracies. (a, b) Acoustic properties of advertisement calls of (a) golden rocket frogs and (b) Kai rocket frogs represented in two dimensions by the first two discriminant functions from the linear discriminant analyses. Each plot represents the calls of five randomly selected individuals. Different colors represent different individuals, larger points outlined in black represent the centroids of individuals’ points, and ellipses represent 95% confidence intervals around the centroids. Note: the same colors on different plots do not necessarily represent the same individuals. (c) Mean (points) and range (vertical lines) of the success with which calls were classified to the correct individual from linear discriminant analyses of 100 randomly selected groups of five individuals for each species. The dotted horizontal line represents the classification success expected by chance.

### (b) Identity reception hypothesis

Males of both species initially responded with aggressive calls and phonotaxis to broadcasts of calls simulating a new neighbor, but this aggression gradually decreased with repeated stimulation during the habituation phase (Fig. 3, Fig. S3). Compared with the first block of the habituation phase, aggression during the last block of the habituation phase was significantly lower for both golden rocket frogs (*V* = 207, *p* < 0.001) and Kai rocket frogs (*V* = 205, *p* < 0.001). Additionally, the aggression of both species decreased at similar rates, as measured by the time required to reach the habituation criteria (golden rocket frogs: mean = 54 min, range = 24 to 96 min; Kai rocket frogs: mean = 65 min, range = 20 to 168 min; *W* = 232, *p* = 0.39). There was a positive relationship between nearest neighbor distance and habituation time for Kai rocket frogs (ρ = 0.67, *p* = 0.001, Fig. S4), but not for golden rocket frogs (ρ = −0.22, *p* = 0.39, Fig. S4). Finally, golden rocket frogs, but not Kai rocket frogs, showed renewed aggression when they heard calls simulating a different male during the discrimination phase. When comparing the last block of the habituation phase with the block of the discrimination phase, there was a significant interaction between treatment and block for golden rocket frogs (*X^2^* = 4.30, *p* = 0.038, Fig. 3a) indicating a greater recovery of aggression in the different frog experimental treatment than the same frog control treatment in this species. Separate tests within treatments revealed a significant increase in aggression for the different frog treatment (*X^2^* = 6.59, *p* = 0.010, Fig. 3b) but not for the same frog treatment (*X^2^* = 3.50, *p* = 0.069, Fig. 3b). In contrast, for Kai rocket frogs, there was no interaction between treatment and block (*X^2^* = 2.39, *p* = 0.122, Fig. 3a) and separate tests revealed no increase in aggression during the discrimination phase for either the different frog treatment (*X^2^* = 0.01, *p* = 0.923, Fig. 3b) or the same frog treatment (*X^2^* = 3.52, *p* = 0.061, Fig 3b).

**Figure 3.**
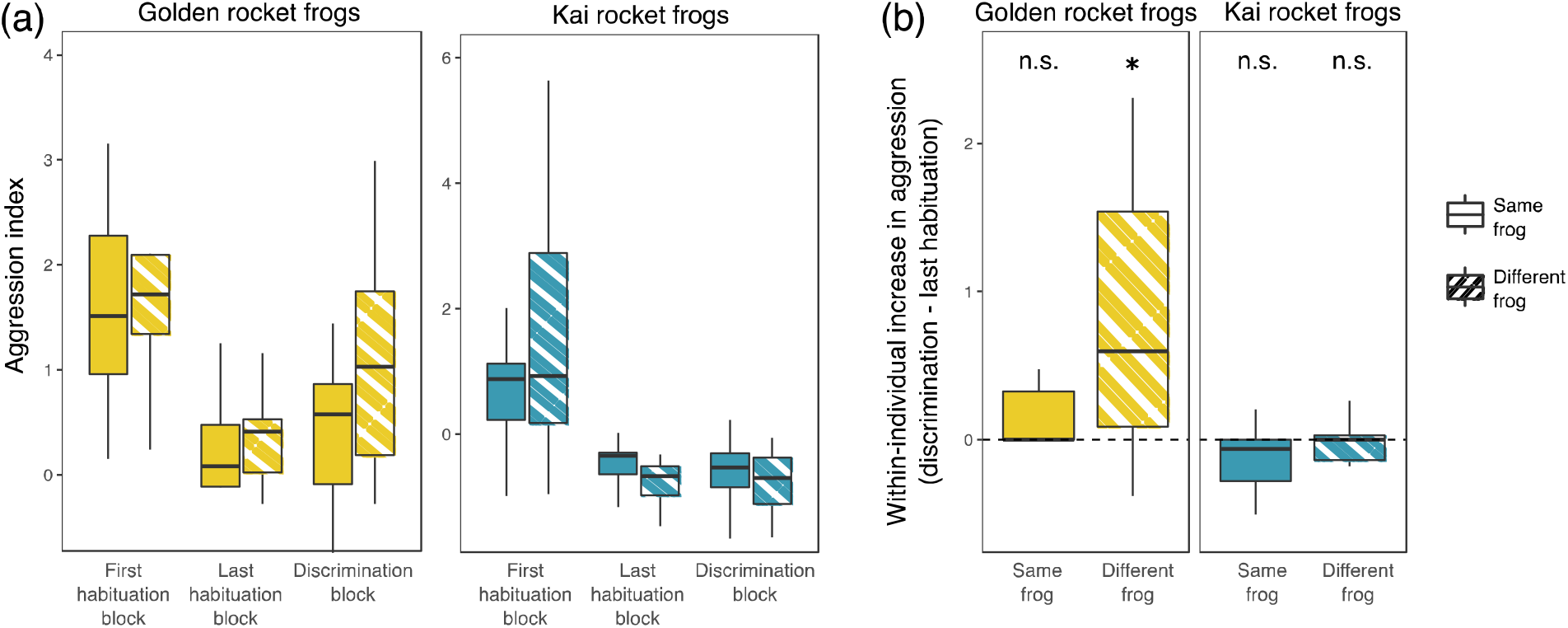
Species difference in the specificity of learned social categories. (a) Boxplots showing the aggression index of subjects during the habituation and discrimination phases of the experiment. Because subjects took different amounts of time to reach habituation criteria, only the first and last blocks of the habituation phase are shown. During the discrimination phase, subjects heard either a different set of calls recorded from the same male (“Same frog,” solid boxes) or calls recorded from a different male (“Different frog,” striped boxes). (b) Boxplots showing the within-individual increase in aggression index between the last block of the habituation phase and the discrimination phase. Asterisk indicates a statistically significant (p < 0.05) increase in aggression, while “n.s.” indicates not significant. (a, b) Horizontal lines show medians, boxes show interquartile ranges, and whiskers extend to the range but no farther than 1.5 times the interquartile range.

## 4. Discussion

### (a) Advertisement calls have not evolved to signal identity

Our results do not strongly support the identity signaling hypothesis: the advertisement calls of both golden rocket frogs and Kai rocket frogs are individually distinctive and convey similar amounts of identity information. The calls of both species show similar patterns of within and between individual variation, with the temporal properties of pulses and the dominant frequency of the call showing the highest potentials for individual coding. These properties would thus be the most useful for identifying individuals in both species. The relatively higher number of pulses in golden rocket frog calls (3 versus 2 on average) contributed to the slightly higher estimate of *H_s_* for this species, and one potential interpretation of this result is that the evolution of calls with additional pulses represents an adaptation to increase the identity information in calls. However, the number of pulses per call for golden rocket frogs vary considerably; even within a bout of calling a male might produce calls with anywhere from 1 to 6 pulses. Moreover, our linear discriminant analysis showed that the difference in *H_s_* between species was negligible because individual differences in the calls of both species can be statistically discriminated with similar accuracy. In groups of five individuals, for example, the calls of both species can be discriminated, on average, with 94% accuracy, which should be more than sufficient for territory owners to make binary discriminations between neighbors and strangers (Fig. 2). Thus, any additional identity information conveyed by having additional pulses in golden rocket frogs is probably not necessary for a functional social recognition system in this species.

Based on our acoustic analyses, we conclude that the evolution of neighbor recognition and the dear enemy effect in golden rocket frogs has not been associated with adaptations in signal production that function to increase acoustic identity information. Likewise, our results confirm that the absence of behavioral discrimination between neighbors and strangers in Kai rocket frogs is unlikely due to limited identity information in their signals. We suggest the advertisement calls of both golden and Kai rocket frogs provide identity *cues* – traits that are individually distinctive but have not evolved to signal identity. What we propose is similar to the way human fingerprints are reliable identifiers of individuals but could not have evolved for that purpose. This suggestion is in line with other studies on frogs showing that many species have individually distinctive calls (reviewed in [18]), including those that do not discriminate between the calls of neighbors and strangers [21,22].

Our lack of support for the identity signaling hypothesis provides an important counterexample to the growing number of studies demonstrating that signals in taxa as varied as swallows [12], paper wasps [37], house mice [38], and humans [39] have evolved to signal identity by providing individual ‘signatures’ [7]. Further, comparisons across species or populations of bats [40], marmots [41], and penguins [42] show patterns consistent with the idea that greater signal identity information is favored in social environments where social recognition is potentially more important. In contrast to these studies, we did not find robust evidence for the evolution of identity signals in golden rocket frogs, despite the expected benefits of reduced aggression from neighbors that would come from being recognized. One possible explanation for the lack of evolved identity signals is that there was sufficient identity information to enable discrimination between neighbors and strangers in the common ancestors of golden and Kai rocket frogs, and thus there was little benefit to increasing identity information beyond this ancestral state. This interpretation is consistent with a coevolutionary model of social recognition showing that selection to increase identity information in signals is relatively weak when existing identity information is already close to the optimum for that recognition system [43].

### (b) Adaptations of identity reception enable social recognition

Our results support the identity reception hypothesis: golden rocket frogs, but not Kai rocket frogs, exhibited habituation of aggression to advertisement calls that was specific to the identity of the stimulus, showing renewed aggression when they subsequently heard calls from a different male during the discrimination phase. Territorial males of both species initially responded with aggression to playbacks of advertisement calls simulating the arrival of a new neighbor, but the aggression gradually decreased with repeated stimulation in a manner consistent with learning through habituation [18,25]. This result is important because it establishes that the decision rule for whether to respond aggressively to a perceived threat is plastic and can be modified by experience in both species. Habituation of aggression appears to be a widespread form of behavioral plasticity among frogs by which males modulate their aggressiveness to accommodate changes in their local social environment [44–46], The key finding from the present study is that only golden rocket frogs exhibited reduced aggression that was specific to the identity of the simulated neighbor. This pattern of learning and discrimination would contribute to allowing males to avoid unnecessary aggression with established neighbors while still allowing them to respond aggressively to unfamiliar strangers [18]. Strikingly similar neighbor-specific habituation of aggression has been demonstrated in North American bullfrogs [6,26,47], a species that also displays a vocally-mediated dear enemy effect [48]. In contrast to golden rocket frogs, Kai rocket frogs did not respond aggressively when presented with the calls of a different male during the discrimination phase, suggesting that their habituation of aggression was not specific to the identity of the simulated neighbor, but rather generalized to the calls of conspecific males. Additional evidence of this generalized habituation comes from the observation that males with nearby neighbors habituated faster to our playbacks than males with more distant neighbors (Fig. S4), suggesting that previous experience with nearby conspecific males caused a general reduction in aggression of Kai rocket frogs. Compared with Kai rocket frogs, golden rocket frogs defend stable reproductive resources and have higher density territories that are defended for longer periods of time [17]; this social environment has presumably selected for the ability to habituate to individual neighbors’ calls, allowing males to avoid repeated but unnecessary aggression with neighbors. Overall, our results are consistent with the hypothesis that an ancestral mechanism of plasticity in aggression that is common among frogs has been evolutionarily adapted through natural selection to become specific to the calls of individual males in golden rocket frogs, thereby enabling social recognition.

Our empirical support for the identity reception hypothesis offers an important and rare example of receivers responding to selection in the context of social recognition. Compared to other components of recognition systems, the perceptual and cognitive basis of social recognition is an aspect of receiver psychology [49] that has received less attention and largely remains a “black box” [10,50], but see [11,13,26,47,51]. Indeed, the combination of perception, learning, and memory that collectively constitutes a receiver’s “psychological landscape” [49] is often implicitly assumed to be a relatively static feature of nervous systems that acts as a source of selection on signals [50]. In contrast, our discovery of a species difference in the specificity of learned social categories highlights how a receiver’s psychological landscape itself can also be a target of selection under conditions favoring social recognition.

### (c) Conclusions

Results from this study indicate that the evolution of social recognition in golden rocket frogs has not been associated with an increase in identity information in signals. Many animal signals vary among individuals, but, as our study highlights, the existence of individual variation does not necessarily mean that signals have evolved to signal identity. Comparative studies such as ours are needed to test the identity signaling hypothesis. Our results instead suggest that mechanisms in receivers that are responsible for determining the specificity of learned social categories have been a key target of selection in the evolution of a social recognition system. Receiver perception and cognition may be more evolutionarily labile than previously appreciated, and comparative studies such as ours highlight the diversity of social cognition, even among closely related species. Future research exploiting such diversity will be needed to reveal in more detail the proximate mechanisms underlying social recognition and how those mechanisms respond to selection.

## Ethics

Research was approved under UMN IACUC Protocol #1701-34456A. Research permits were provided by the Guyana EPA (060214 BR 018 and 040717 BR 004) and permission was granted by the Guyana Protected Areas Commission.

## Data accessibility

Data will be deposited in a public repository upon publication.

## Author contributions

J.P.T. and M.A.B. conceived the study and acquired funding. J.P.T. and Z.K.L. collected the data. J.P.T analyzed the data. J.P.T. and M.A.B. wrote the manuscript with contributions from Z.K.L.

## Acknowledgements

We thank Godfrey Bourne and Beth Pettitt for advice and logistical assistance, and Maxwell Basil and Thomas John for field assistance.

## References

1. Colgan, P. W. 1983 Comparative social recognition. Wiley.

2. Bee, M. A. 2006 Individual recognition in animal species. In The Encyclopedia of Language and Linguistics: Volume 2 (ed N. Naguib), pp. 617–626. London: Elsevier Science.

3. Sidtis, D. V. L. & Zäske, R. 2021 Who We Are. In The Handbook of Speech Perception, pp. 365–397. Wiley. (doi:10.1002/9781119184096.ch14)

4. Wiley, R. H. 2013 Specificity and multiplicity in the recognition of individuals: implications for the evolution of social behaviour. Biol. Rev. 88, 179–95. (doi:10.1111/j.1469-185X.2012.00246.x)

5. Tibbetts, E. A., Pardo-Sanchez, J., Ramirez-Matias, J. & Avarguès-Weber, A. 2021 Individual recognition is associated with holistic face processing in Polistes paper wasps in a species-specific way. Proc. R. Soc. B Biol. Sci. 288, 20203010. (doi:10.1098/rspb.2020.3010)

6. Bee, M. A. & Gerhardt, H. C. 2002 Individual voice recognition in a territorial frog (*Rana catesbeiana*). Proc. R. Soc. London B Biol. Sci. 269, 1443–1448. (doi:10.1098/rspb.2002.2041)

7. Beecher, M. D. 1989 Signalling systems for individual recognition: an information theory approach. Anim. Behav. 38, 248–261. (doi:10.1016/S0003-3472(89)80087-9)

8. Tibbetts, E. A. & Dale, J. 2007 Individual recognition: it is good to be different. Trends Ecol. Evol. 22, 529–37. (doi:10.1016/j.tree.2007.09.001)

9. Sherman, P. W., Reeve, H. K. & Pfennig, D. W. 1997 Recognition systems. In Behavioral ecology: An evolutionary approach (eds J. R. Krebs & N. B. Davies), pp. 69–96. Oxford: Blackwell Science Ltd.

10. Yorzinski, J. L. 2017 The cognitive basis of individual recognition. Curr. Opin. Behav. Sci. 16, 53–57. (doi:10.1016/j.cobeha.2017.03.009)

11. Loesche, P., Stoddard, P. K., Higgins, B. J. & Beecher, M. D. 1991 Signature versus perceptual adaptations for individual vocal recognition in swallows. Behaviour 118, 15–25.

12. Medvin, M. B., Stoddard, P. K. & Beecher, M. D. 1993 Signals for parent-offspring recognition: a comparative analysis of begging calls of cliff swallows and barn swallows. Anim. Behav. 45, 841–850.

13. Sheehan, M. J. & Tibbetts, E. A. 2011 Specialized face learning is associated with individual recognition in paper wasps. Science. 334, 1272–1275. (doi:10.1126/science.1211334)

14. Temeles, E. 1994 The role of neighbours in territorial systems: when are they ‘dear enemies’? Anim. Behav. 47, 339–350.

15. Stoddard, P. K. 1996 Vocal recognition of neighbors by territorial passerines. In Ecology and Evolution of Acoustic Communication in Birds (eds D. E. Kroodsma & E. H. Miller), pp. 356–34. Ithaca, NY: Cornell University Press.

16. Wilson, E. O. 1975 Sociobiology: The New Synthesis. Cambridge, MA: Harvard University Press.

17. Tumulty, J. P. & Bee, M. A. 2021 Ecological and social drivers of neighbor recognition and the dear enemy effect in a poison frog. Behav. Ecol. 32, 138–150. (doi:10.1093/beheco/araa113)

18. Bee, M. A., Reichert, M. S. & Tumulty, J. 2016 Assessment and recognition of competitive rivals in anuran amphibians. Adv. Study Behav. 48, 161–249. (doi:10.1016/bs.asb.2016.01.001)

19. Bourne, G. R., Collins, A., Holder, A. & McCarthy, C. 2001 Vocal communication and reproductive behavior of the frog *Colostethus beebei* in Guyana. J. Herpetol. 35, 272–281.

20. Pettitt, B. A., Bourne, G. R. & Bee, M. A. 2013 Advertisement call variation in the golden rocket frog (*Anomaloglossus beebei*): evidence for individual distinctiveness. Ethology 119, 244–256.

21. Bee, M. A. 2003 A test of the ‘dear enemy effect’ in the strawberry dart-poison frog (*Dendrobates pumilio*). Behav. Ecol. Sociobiol. 54, 601–610. (doi:10.1007/s00265-003-0657-5)

22. Tumulty, J. P., Pašukonis, A., Ringler, M., Forester, J. D., Hödl, W. & Bee, M. A. 2018 Brilliant-thighed poison frogs do not use acoustic identity information to treat territorial neighbours as dear enemies. Anim. Behav. 141, 203–220. (doi:10.1016/j.anbehav.2018.05.008)

23. Peeke, H. V. & Peeke, S. C. 1973 Habituation in fish with special reference to intraspecific aggressive behavior. In Habituation: Behavioral studies, pp. 59–83.

24. Petrinovich, L. 1984 A two-factor dual-process theory of habituation and sensitization. In Habituation, sensitization, and behaviour (eds H. V. S. Peeke & L. Petrinovich), New York: Academic Press.

25. Shettleworth, S. J. 2009 Cognition, evolution, and behavior. New York, NY: Oxford University Press.

26. Bee, M. A. & Gerhardt, H. C. 2001 Habituation as a mechanism of reduced aggression between neighboring territorial male bullfrogs (*Rana catesbeiana*). J. Comp. Psychol. 115, 68–82. (doi:10.1037//073S-7036.115.1.68)

27. The Herpetological Animal Care and Use Committee (HACC) of American Society of Ichthyologists and Herpetologists 2004 Guidelines for use of live amphibians and reptiles in field and laboratory research, 2nd ed.

28. Kok, P. J., Sambhu, H., Roopsind, I., Lenglet, G. L. & Bourne, G. R. 2006 A new species of *Colostethus* (Anura: Dendrobatidae) with maternal care from Kaieteur National Park, Guyana. Zootaxa 1238, 35–61.

29. Pettitt, B. A., Bourne, G. R. & Bee, M. A. 2018 Predictors and benefits of microhabitat selection for offspring deposition in golden rocket frogs. Biotropica 50, 919–928. (doi:10.1111/btp.12609)

30. Pettitt, B. A., Bourne, G. R. & Bee, M. A. 2020 Females prefer the calls of better fathers in a Neotropical frog with biparental care. Behav. Ecol. 31, 152–163. (doi:10.1093/beheco/arz172)

31. Pettitt, B. A., Bourne, G. R. & Bee, M. A. 2012 Quantitative acoustic analysis of the vocal repertoire of the golden rocket frog (*Anomaloglossus beebei*). J. Acoust. Soc. Am. 131, 4811–20. (doi:10.1121/1.4714769)

32. Vacher, J. P. et al. 2017 Cryptic diversity in Amazonian frogs: Integrative taxonomy of the genus *Anomaloglossus* (Amphibia: Anura: Aromobatidae) reveals a unique case of diversification within the Guiana Shield. Mol. Phylogenet. Evol. 112, 158–173. (doi:10.1016/j.ympev.2017.04.017)

33. Bee, M. A., Kozich, C. E., Blackwell, K. J. & Gerhardt, H. C. 2001 Individual variation in advertisement calls of territorial male green frogs, *Rana clamitans:* implications for individual discrimination. Ethology 107, 65–84.

34. Bee, M. A. & Gerhardt, H. C. 2001 Neighbour–stranger discrimination by territorial male bullfrogs (*Rana catesbeiana*): I. Acoustic basis. Anim. Behav. 62, 1129–1140. (doi:10.1006/anbe.2001.1851)

35. Chuang, M.-F., Kam, Y.-C. & Bee, M. A. 2017 Territorial olive frogs display lower aggression towards neighbours than strangers based on individual vocal signatures. Anim. Behav. 123, 217–228. (doi:10.1016/j.anbehav.2016.11.001)

36. Linhart, P., Osiejuk, T. S., Budka, M., Šálek, M., Špinka, M., Policht, R., Syrová, M. & Blumstein, D. T. 2019 Measuring individual identity information in animal signals: Overview and performance of available identity metrics. Methods Ecol. Evol. 10, 1558–1570. (doi:10.1111/2041-210X.13238)

37. Sheehan, M. J. & Tibbetts, E. A. 2010 Selection for individual recognition and the evolution of polymorphic identity signals in *Polistes* paper wasps. J. Evol. Biol. 23, 570–7. (doi:10.1111/j.1420-9101.2009.01923.x)

38. Sheehan, M. J., Lee, V., Corbett-Detig, R., Bi, K., Beynon, R. J., Hurst, J. L. & Nachman, M. W. 2016 Selection on coding and regulatory variation maintains individuality in major urinary protein scent marks in wild mice. PLoS Genet. 12, 1–33. (doi:10.1371/journal.pgen.1005891)

39. Sheehan, M. J. & Nachman, M. W. 2014 Morphological and population genomic evidence that human faces have evolved to signal individual identity. Nat. Commun. 5, 4800. (doi:10.1038/ncomms5800)

40. Wilkinson, G. S. 2003 Social and vocal complexity in bats. In Animal Social Complexity (eds F. B. de Waal & P. L. Tyack), pp. 322–341. Cambridge, MA: Harvard University Press.

41. Pollard, K. A. & Blumstein, D. T. 2011 Social group size predicts the evolution of individuality. Curr. Biol. 21, 413–7. (doi:10.1016/j.cub.2011.01.051)

42. Aubin, T. & Jouventin, P. 2002 How to vocally identify kin in a crowd: the penguin model. Adv. Study Behav. 31, 243–277. (doi:10.1016/S0065-3454(02)80010-9)

43. Miller, S. E., Sheehan, M. J. & Reeve, H. K. 2020 Coevolution of cognitive abilities and identity signals in individual recognition systems: Selection on senders and receivers. Philos. Trans. R. Soc. B Biol. Sci. 375. (doi:10.1098/rstb.2019.0467)

44. Brenowitz, E. A. & Rose, G. J. 1994 Behavioural plasticity mediates aggression in choruses of the Pacific treefrog. Anim. Behav. 47, 633–41.

45. Marshall, V. T., Humfeld, S. C. & Bee, M. A. 2003 Plasticity of aggressive signalling and its evolution in male spring peepers, *Pseudacris crucifer*. Anim. Behav. 65, 1223–1234. (doi:10.1006/anbe.2003.2134)

46. Reichert, M. S. 2010 Aggressive thresholds in *Dendropsophus ebraccatus*: Habituation and sensitization to different call types. Behav. Ecol. Sociobiol. 64, 529–539. (doi:10.1007/s00265-009-0868-5)

47. Bee, M. A. & Gerhardt, H. C. 2001 Neighbour–stranger discrimination by territorial male bullfrogs (*Rana catesbeiana*): II. Perceptual basis. Anim. Behav. 62, 1141–1150. (doi:10.1006/anbe.2001.1852)

48. Davis, M. S. 1987 Acoustically mediated neighbor recognition in the North American bullfrog, *Rana catesbeiana*. Behav. Ecol. Sociobiol. 21, 185–190. (doi:10.1007/BF00303209)

49. Guilford, T. & Dawkins, M. S. 1991 Receiver psychology and the evolution of animal signals. Anim. Behav. 42, 1–14.

50. Miller, C. T. & Bee, M. A. 2012 Receiver psychology turns 20: is it time for a broader approach? Anim. Behav. 83, 331–343. (doi:10.1016/j.anbehav.2011.11.025)

51. Miller, S. E., Legan, A. W., Henshaw, M. T., Ostevik, K. L., Samuk, K., Uy, F. M. K. & Sheehan, M. J. 2020 Evolutionary dynamics of recent selection on cognitive abilities. Proc. Natl. Acad. Sci. U. S. A. 117, 3045–3052. (doi:10.1073/pnas.1918592117)

