## Supplementary material for "Identity signaling, identity reception and the evolution of social recognition in a Neotropical frog"

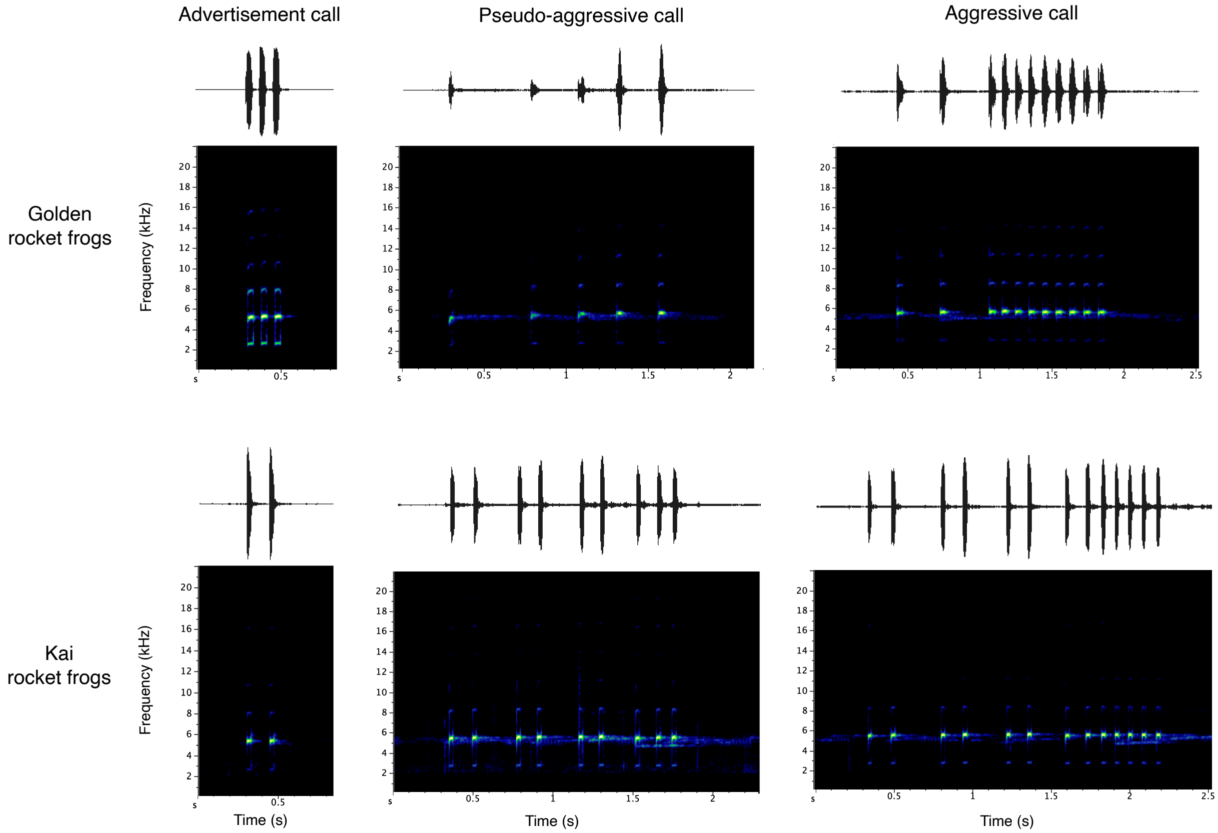


**Figure S1.** Vocal repertoire of golden rocket frogs and Kai rocket frogs. Advertisement calls for golden rocket frog consist of a series of average of 3 pulses (range 1-6) consist of series of 3 pulses (range 1-6) for golden rocket frogs or 2 pulses (range 1-2) for Kai rocket frogs with dominant frequencies in the range of 4.6-5.8 kHz. Aggressive calls are longer and consist of several introductory pulses (in golden rocket frogs) or rapid advertisement calls (in Kai rocket frogs) followed by a long train of pulses with relatively shorter inter-pulse intervals. Because of variation in the presence of introductory pulses given by golden rocket frogs, we classified all calls with at least seven pulses as aggressive calls in this species. Additionally, both species sometimes produce what we refer to as "pseudo-aggressive calls," which consist of the characteristic introductory pulses without a subsequent train of pulses in golden rocket frogs, or several rapid advertisement calls followed by a three-pulse call in Kai rocket frogs.

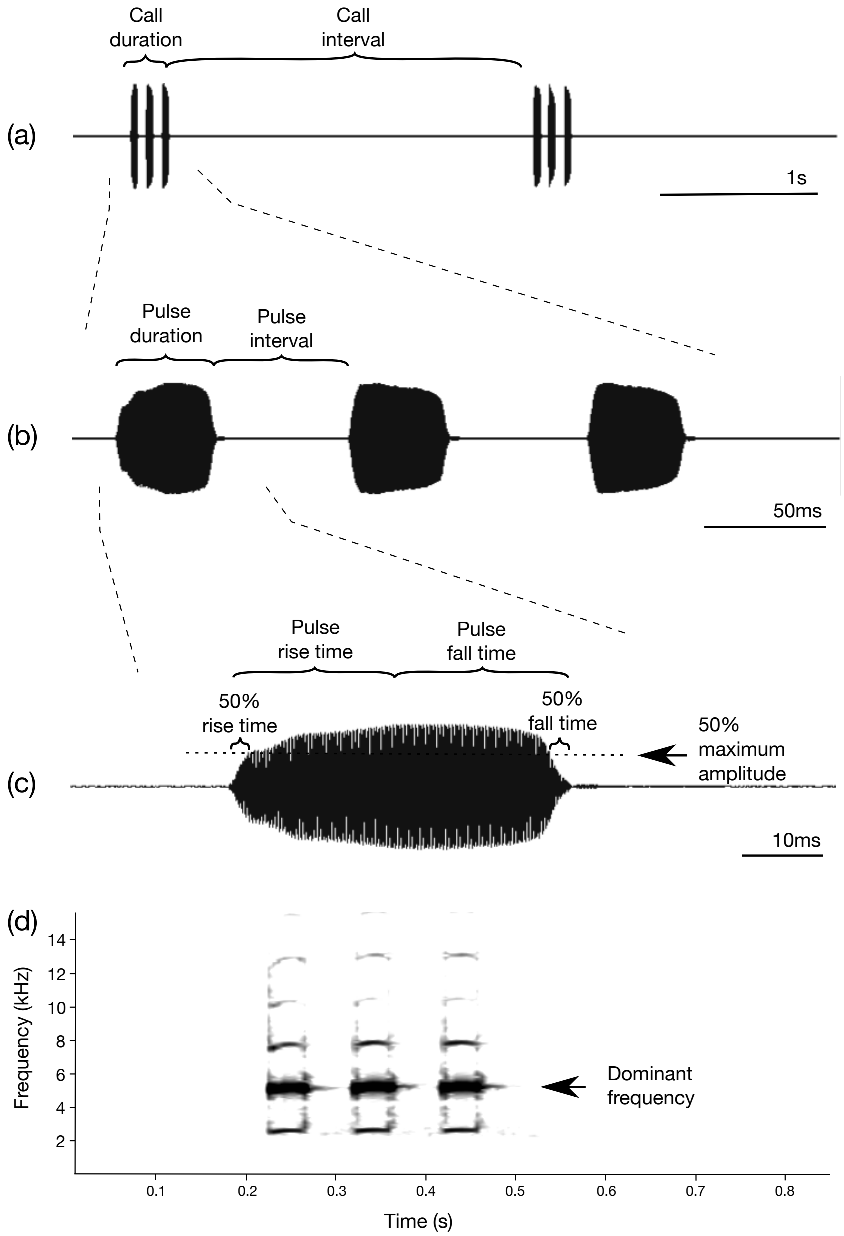


**Figure S2.** Waveforms (a-c) and spectrogram (d) of golden rocket frog advertisement calls showing the acoustic properties that were measured to quantify identity information. We measured call duration as the time from the onset of the first pulse to the offset of the last pulse in a call, and the call interval as the time from the offset of the last pulse in a call to the onset of the first pulse in the subsequent call. Call period was the sum of call duration and call interval. For each pulse in a call, we measured pulse duration as the time between the onset and the offset of the pulse and pulse interval as the time between the offset of the pulse and the onset of the subsequent pulse. The sum of pulse duration and pulse interval was pulse period. Additionally, we measured four temporal properties to describe pulse shape: pulse rise time (onset to maximum amplitude), 50% rise time (onset to 50% of maximum amplitude), fall time (maximum amplitude to offset) and 50% fall time (50% of maximum amplitude to offset). We measured the dominant frequency of each pulse using the max frequency function on the power spectrum of the pulse (1024 point, Hamming window). The same acoustic properties were measured on Kai rocket frog advertisement calls.

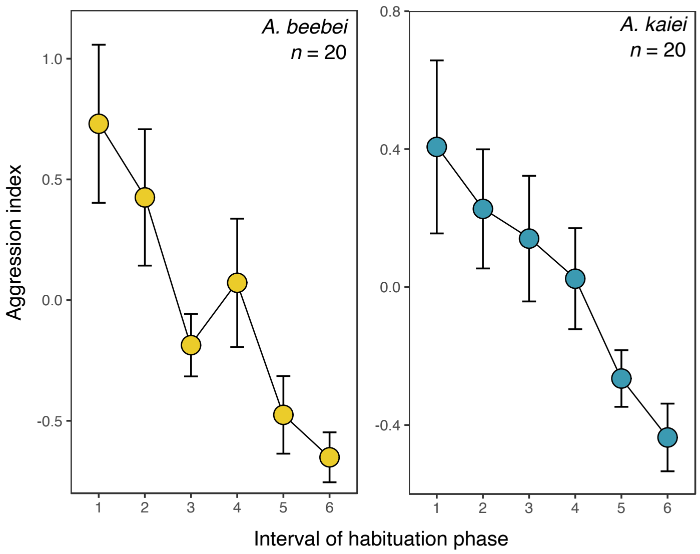


**Figure S3.** The mean (±SE) aggression index of territorial males in response to repeated broadcasts of advertisement call playback during the habituation phase. Subjects took different amounts of time to reach habituation criteria so here we visualize changes in aggression during the habituation phase by partitioning the habituation phase into 6 intervals, each constituting 16.7% of a subject's total time to reach habituation criteria.

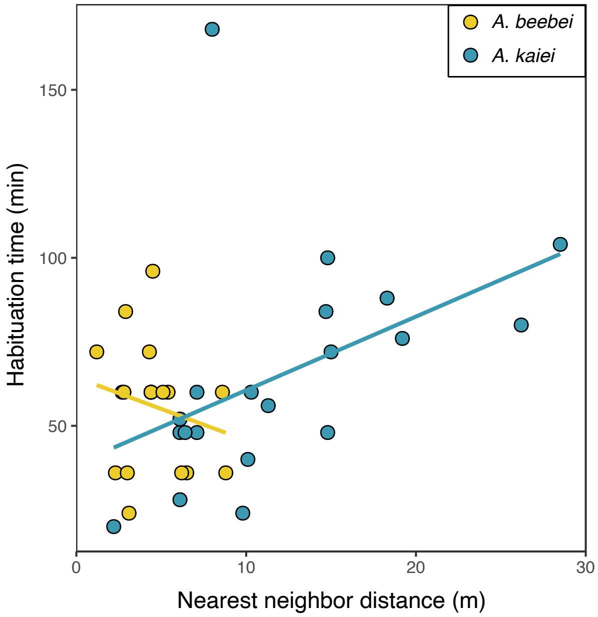


**Figure S4.** The relationship between the time to reach habituation criteria and nearest neighbor distance fit with a regression line for each species. There was a positive relationship between nearest neighbor distance and habituation time for Kai rocket frogs ($\rho$ = 0.67, $p$ = 0.001), but not for golden rocket frogs ($\rho$ = -0.22, $p$ = 0.39).

**Table S1**. Descriptive statistics of within- and among-individual variation in acoustic properties of golden rocket frog advertisement calls, for each pulse separately, including estimates of the potential for individual coding (PIC) and effect sizes associated with individual differences (partial $\eta^2$).

| Call Property | | Mean | SD | $CV_a$ | Mean $CV_w$ (range) | PIC | Partial $\eta^2$ |
|---|---|---|---|---|---|---|---|
| Call temporal properties | Call duration (s) | 0.229 | 0.044 | 19.4 | 16.5 (1.2-28.6) | 1.2 | 0.54 |
| | No. pulses | 3.16 | 0.52 | 16.5 | 12.8 (0-23.3) | 1.3 | 0.57 |
| | Call interval (s) | 2.516 | 0.821 | 32.6 | 39.1 (6.1-198.7) | 0.8 | 0.09 |
| | Call period (s) | 2.744 | 0.809 | 29.5 | 36.8 (5.7-191.1) | 0.8 | 0.08 |
| Pulse duration (ms) | Pulse 1 | 36.2 | 3.4 | 9.4 | 3.9 (1.9-5.8) | 2.4 | 0.84 |
| | Pulse 2 | 36.0 | 3.1 | 8.6 | 3.6 (2-8.9) | 2.4 | 0.82 |
| | Pulse 3 | 35.4 | 3.0 | 8.5 | 4.0 (1.5-8.4) | 2.1 | 0.78 |
| | Pulse 4 | 33.0 | 3.6 | 10.9 | 7.4 (2.1-25.9) | 1.5 | 0.78 |
| Pulse interval (ms) | Pulse 1 | 53.7 | 5.9 | 11.0 | 5.1 (2-7.6) | 2.1 | 0.81 |
| | Pulse 2 | 53.3 | 4.1 | 7.6 | 4.2 (2.3-7.4) | 1.8 | 0.77 |
| | Pulse 3 | 55.5 | 4.3 | 7.8 | 6.1 (2.9-9.1) | 1.3 | 0.43 |
| Pulse period (ms) | Pulse 1 | 90.0 | 7.0 | 7.7 | 2.8 (1.7-4.4) | 2.8 | 0.88 |
| | Pulse 2 | 89.4 | 5.1 | 5.7 | 2.4 (1.3-4.6) | 2.4 | 0.86 |
| | Pulse 3 | 91.3 | 5.0 | 5.4 | 3.4 (1.7-6.7) | 1.6 | 0.63 |
| Pulse rise time (ms) | Pulse 1 | 18.0 | 6.2 | 34.1 | 37.5 (18.9-66.0) | 0.9 | 0.46 |
| | Pulse 2 | 15.0 | 4.2 | 28.2 | 45.0 (22.8-110.9) | 0.6 | 0.27 |
| | Pulse 3 | 14.2 | 4.0 | 27.8 | 43.5 (20.7-89.0) | 0.6 | 0.31 |
| | Pulse 4 | 13.3 | 3.9 | 28.9 | 31.8 (4.1-74.6) | 0.9 | 0.35 |
| Pulse fall time (ms) | Pulse 1 | 18.2 | 5.8 | 32.1 | 40.0 (12.3-69.3) | 0.8 | 0.42 |
| | Pulse 2 | 21.1 | 5.0 | 23.8 | 31.9 (13.0-57.8) | 0.7 | 0.35 |
| | Pulse 3 | 21.1 | 5.0 | 23.7 | 29.9 (9.4-84.6) | 0.8 | 0.40 |
| | Pulse 4 | 19.7 | 5.7 | 29.0 | 26.5 (5.1-62.0) | 1.1 | 0.44 |
| Pulse 50% rise time (ms) | Pulse 1 | 4.3 | 3.3 | 75.6 | 50.7 (21.3-124.3) | 1.5 | 0.48 |
| | Pulse 2 | 2.9 | 1.5 | 51.1 | 44.2 (0-125.6) | 1.2 | 0.36 |
| | Pulse 3 | 2.6 | 1.2 | 45.1 | 32.4 (0-57.0) | 1.4 | 0.54 |
| | Pulse 4 | 3.2 | 1.5 | 48.7 | 35.6 (0-86.5) | 1.4 | 0.54 |
| Pulse 50% fall time (ms) | Pulse 1 | 3.6 | 1.6 | 44.5 | 74.4 (30.9-152.5) | 0.6 | 0.17 |
| | Pulse 2 | 3.6 | 1.4 | 37.8 | 65.1 (19.2-163.2) | 0.6 | 0.16 |
| | Pulse 3 | 4.0 | 1.8 | 45.8 | 65.2 (19.4-158.0) | 0.7 | 0.20 |
| | Pulse 4 | 4.9 | 5.2 | 106.1 | 47.5 (0-81.7) | 2.2 | 0.40 |
| Dominant frequency (Hz) | Pulse 1 | 5257 | 216 | 4.1 | 2.4 (0.6-14.8) | 1.8 | 0.50 |
| | Pulse 2 | 5342 | 248 | 4.6 | 1.8 (0.3-12.0) | 2.6 | 0.65 |
| | Pulse 3 | 5360 | 256 | 4.8 | 1.2 (0.5-2.3) | 4.0 | 0.85 |
| | Pulse 4 | 5300 | 175 | 3.3 | 1.2 (0-2.4) | 2.7 | 0.85 |

**Table S2.** Descriptive statistics of within- and among-individual variation in acoustic properties of Kai rocket frog advertisement calls, for each pulse separately, including estimates of the potential for individual coding (PIC) and effect sizes associated with individual differences (partial $\eta^2$).

| Call property | | Mean | SD | $CV_a$ | Mean $CV_w$ (range) | PIC | Partial $\eta^2$ |
|---|---|---|---|---|---|---|---|
| Call temporal properties | Call duration (s) | 0.154 | 0.017 | 10.8 | 11.4 (0.9-59.9) | 0.9 | 0.31 |
| | No. pulses | 1.958 | 0.094 | 4.8 | 5.3 (0-29.7) | 0.9 | 0.20 |
| | Call interval (s) | 0.855 | 0.369 | 43.1 | 40.1 (11.7-154.1) | 1.1 | 0.22 |
| | Call period (s) | 1.009 | 0.370 | 36.7 | 34.5 (9.4-140.8) | 1.1 | 0.22 |
| Pulse duration (ms) | Pulse 1 | 24.0 | 3.4 | 14.0 | 6.7 (2.7-13.6) | 2.1 | 0.79 |
| | Pulse 2 | 26.5 | 2.4 | 9.0 | 4.6 (2.3-10.7) | 1.9 | 0.79 |
| Pulse period (ms) | Pulse 1 | 134.8 | 7.7 | 5.7 | 2.3 (1.0-3.8) | 2.5 | 0.65 |
| Pulse interval (ms) | Pulse 1 | 110.5 | 7.1 | 6.4 | 3.1 (1.8-4.7) | 2.1 | 0.59 |
| Pulse rise time (ms) | Pulse 1 | 9.1 | 4.4 | 48.5 | 41.3 (16.6-66.5) | 1.2 | 0.58 |
| | Pulse 2 | 9.4 | 3.9 | 41.1 | 40.7 (6.0-58.0) | 1.0 | 0.50 |
| Pulse fall time (ms) | Pulse 1 | 14.9 | 3.8 | 25.7 | 29.3 (13.8-72.0) | 0.9 | 0.46 |
| | Pulse 2 | 17.2 | 4.0 | 23.5 | 25.8 (5.3-49.3) | 0.9 | 0.46 |
| Pulse 50% rise time (ms) | Pulse 1 | 2.1 | 1.0 | 46.2 | 38.0 (0-104.0) | 1.2 | 0.44 |
| | Pulse 2 | 1.9 | 0.9 | 48.3 | 28.5 (0-47.9) | 1.7 | 0.61 |
| Pulse 50% fall time (ms) | Pulse 1 | 3.0 | 2.3 | 75.9 | 53.5 (21.7-94.5) | 1.4 | 0.55 |
| | Pulse 2 | 2.8 | 1.5 | 52.5 | 51.7 (28.2-89.0) | 1.0 | 0.57 |
| Dominant frequency (Hz) | Pulse 1 | 5179 | 162 | 3.1 | 1.5 (0.6-4.1) | 2.2 | 0.79 |
| | Pulse 2 | 5201 | 179 | 3.4 | 1.2 (0.5-3.3) | 3.0 | 0.83 |

**Table S3.** Acoustic identity information and classification success using different inclusion criteria for principal components (PCs).

| | No. PCs | Beecher's $H_s$ | Classification success for full data set (%) | $\chi^2$ | p-value | Classification success for groups of 5 individuals (%) mean | min | max |
|---|---|---|---|---|---|---|---|---|
| *PCs that explain > 99% of variation* | | | | | | | | |
| Golden rocket frogs | 17 | 5.09 | 86.29 | | | 94.3 | 84.62 | 100 |
| Kai rocket frogs* | 11 | 4.57 | 84.07 | 0.21 | 0.646 | 93.9 | 80.43 | 100 |
| *PCs that explain 95% of variation* | | | | | | | | |
| Golden rocket frogs | 13 | 4.86 | 84.86 | | | 95.2 | 87.32 | 100 |
| Kai rocket frogs | 9 | 4.21 | 79.63 | 0.93 | 0.335 | 93.4 | 78.26 | 99 |
| *PCs with SD > 1* | | | | | | | | |
| Golden rocket frogs | 9 | 4.36 | 80.86 | | | 93.6 | 81.97 | 100 |
| Kai rocket frogs | 6 | 3.87 | 77.02 | 0.52 | 0.472 | 92.2 | 77.66 | 98.95 |
| *PCs from first two pulses (>99% of variation)* | | | | | | | | |
| Golden rocket frogs | 12 | 4.31 | 76.63 | | | 92.9 | 80.61 | 100 |
| Kai rocket frogs* | 11 | 4.57 | 84.07 | 2.27 | 0.124 | 93.9 | 80.43 | 100 |

*Kai rocket frog calls have two pulses, so the same data are presented twice for reference, once for the first analysis (as presented in the main text) and again for the analysis that restricts the golden rocket frog data set to properties of the first two pulses.

**Table S4.** Loadings of measured golden rocket frog call properties on principal components and the variance explained by each component.

| | | PCA Factor | | | | | | | | | | | | | | | | |
|---|---|---|---|---|---|---|---|---|---|---|---|---|---|---|---|---|---|---|
| Call property | | 1 | 2 | 3 | 4 | 5 | 6 | 7 | 8 | 9 | 10 | 11 | 12 | 13 | 14 | 15 | 16 | 17 |
| Call temporal properties | Call duration | -0.18 | -0.06 | 0.16 | -0.21 | -0.22 | **0.45** | 0.24 | **0.30** | 0.06 | 0.05 | -0.02 | -0.04 | -0.08 | -0.03 | 0.00 | -0.02 | 0.00 |
| | No. pulses | -0.15 | 0.04 | **0.26** | -0.17 | -0.22 | **0.42** | 0.21 | **0.29** | 0.05 | 0.07 | -0.05 | -0.04 | -0.07 | -0.03 | 0.01 | -0.05 | -0.04 |
| | Call interval | 0.04 | 0.00 | -0.15 | **0.26** | **-0.61** | -0.12 | 0.01 | 0.03 | -0.10 | -0.02 | 0.00 | 0.00 | 0.02 | 0.01 | -0.01 | -0.01 | 0.01 |
| | Call period | 0.04 | 0.00 | -0.15 | **0.26** | **-0.62** | -0.12 | 0.01 | 0.04 | -0.10 | -0.02 | -0.01 | 0.00 | 0.02 | 0.00 | -0.01 | -0.01 | 0.01 |
| Pulse duration | Pulse 1 | -0.11 | **-0.37** | -0.04 | -0.18 | -0.03 | -0.21 | 0.19 | -0.11 | 0.00 | -0.05 | 0.02 | -0.14 | 0.06 | -0.15 | 0.05 | **-0.68** | -0.08 |
| | Pulse 2 | -0.12 | **-0.37** | 0.02 | -0.17 | -0.04 | -0.15 | 0.17 | -0.05 | -0.13 | -0.17 | 0.05 | 0.22 | -0.04 | -0.12 | -0.10 | **0.40** | **-0.49** |
| | Pulse 3 | -0.14 | **-0.36** | 0.10 | -0.15 | -0.04 | -0.10 | 0.14 | -0.04 | -0.15 | -0.23 | 0.06 | 0.09 | 0.03 | **0.31** | 0.11 | 0.19 | **0.68** |
| Pulse interval | Pulse 1 | 0.05 | -0.01 | **-0.47** | -0.03 | 0.03 | **0.25** | -0.09 | -0.05 | 0.08 | -0.01 | 0.14 | -0.05 | **-0.48** | -0.06 | 0.01 | 0.24 | 0.11 |
| | Pulse 2 | -0.06 | -0.08 | **-0.43** | 0.09 | 0.09 | 0.24 | -0.05 | 0.13 | 0.17 | 0.14 | -0.21 | -0.01 | **0.47** | 0.18 | 0.01 | -0.08 | 0.11 |
| Pulse period | Pulse 1 | -0.02 | -0.21 | **-0.43** | -0.12 | 0.01 | 0.10 | 0.03 | -0.10 | 0.06 | -0.04 | 0.14 | -0.13 | **-0.38** | -0.14 | 0.04 | -0.17 | 0.05 |
| | Pulse 2 | -0.12 | **-0.29** | **-0.35** | -0.03 | 0.05 | 0.11 | 0.06 | 0.08 | 0.07 | 0.01 | -0.15 | 0.12 | **0.37** | 0.08 | -0.05 | 0.18 | -0.20 |
| Pulse rise time | Pulse 1 | **-0.33** | -0.06 | -0.01 | -0.08 | -0.03 | 0.13 | -0.24 | -0.21 | **-0.38** | 0.20 | 0.12 | 0.16 | 0.05 | -0.05 | 0.03 | -0.24 | -0.03 |
| | Pulse 2 | **-0.34** | 0.14 | -0.11 | -0.04 | 0.04 | -0.16 | 0.09 | 0.18 | -0.06 | -0.21 | **0.41** | -0.24 | 0.17 | -0.07 | 0.00 | 0.16 | -0.13 |
| | Pulse 3 | **-0.27** | 0.18 | -0.15 | -0.15 | 0.06 | **-0.26** | 0.15 | 0.18 | -0.20 | -0.09 | **-0.41** | 0.14 | -0.19 | 0.08 | -0.06 | -0.01 | 0.19 |
| Pulse fall time | Pulse 1 | **0.29** | -0.10 | -0.01 | 0.00 | 0.02 | -0.22 | **0.32** | 0.17 | **0.39** | -0.22 | -0.11 | -0.22 | -0.03 | -0.02 | -0.01 | -0.05 | -0.01 |
| | Pulse 2 | **0.27** | **-0.29** | 0.11 | -0.03 | -0.06 | 0.09 | -0.01 | -0.19 | 0.01 | 0.13 | **-0.37** | **0.32** | -0.17 | 0.02 | -0.04 | 0.02 | -0.09 |
| | Pulse 3 | 0.18 | **-0.32** | 0.18 | 0.07 | -0.07 | 0.19 | -0.07 | -0.18 | 0.12 | -0.02 | **0.40** | -0.08 | 0.19 | 0.06 | 0.10 | 0.09 | 0.12 |
| Pulse 50% rise time | Pulse 1 | **-0.25** | -0.07 | 0.09 | -0.14 | -0.13 | -0.11 | -0.03 | **-0.38** | 0.11 | **0.37** | **-0.30** | **-0.61** | 0.03 | -0.10 | -0.13 | 0.29 | 0.07 |
| | Pulse 2 | -0.20 | 0.15 | 0.02 | -0.21 | -0.18 | -0.03 | -0.15 | -0.22 | **0.48** | -0.14 | 0.16 | **0.25** | -0.02 | **0.27** | **-0.59** | -0.14 | 0.01 |
| | Pulse 3 | -0.18 | 0.19 | 0.00 | -0.25 | -0.19 | -0.06 | -0.16 | -0.20 | **0.40** | -0.16 | -0.11 | 0.23 | 0.08 | -0.17 | **0.69** | 0.07 | -0.03 |
| Pulse 50% fall time | Pulse 1 | 0.04 | -0.07 | -0.05 | -0.17 | -0.02 | **-0.37** | 0.08 | **0.30** | 0.18 | **0.73** | **0.30** | 0.23 | -0.06 | 0.05 | 0.09 | 0.05 | 0.07 |
| | Pulse 2 | 0.04 | -0.20 | 0.07 | -0.11 | -0.03 | -0.08 | **-0.52** | **0.37** | 0.06 | -0.11 | -0.09 | 0.04 | 0.10 | **-0.61** | -0.24 | 0.05 | **0.25** |
| | Pulse 3 | 0.00 | -0.19 | 0.06 | -0.15 | -0.06 | -0.09 | **-0.54** | **0.31** | -0.02 | -0.08 | -0.08 | **-0.28** | -0.19 | **0.55** | 0.17 | -0.05 | **-0.28** |
| Dominant frequency | Pulse 1 | 0.30 | 0.14 | -0.14 | **-0.34** | -0.08 | 0.07 | -0.03 | -0.03 | -0.18 | -0.03 | 0.03 | -0.01 | 0.17 | -0.02 | 0.06 | -0.06 | 0.01 |
| | Pulse 2 | **0.28** | 0.14 | -0.10 | **-0.41** | -0.11 | 0.00 | 0.03 | -0.05 | -0.18 | 0.01 | 0.02 | -0.05 | 0.15 | 0.04 | -0.09 | 0.07 | -0.03 |
| | Pulse 3 | **0.28** | 0.14 | -0.07 | **-0.42** | -0.10 | -0.02 | 0.03 | -0.05 | -0.16 | -0.03 | 0.05 | -0.08 | 0.11 | 0.03 | -0.07 | 0.04 | 0.03 |
| Variance | | 5.08 | 4.54 | 3.41 | 2.23 | 1.9 | 1.65 | 1.54 | 1.31 | 1.14 | 0.82 | 0.49 | 0.45 | 0.39 | 0.31 | 0.27 | 0.19 | 0.15 |
| Proportion of Variance | | 0.2 | 0.17 | 0.13 | 0.09 | 0.07 | 0.06 | 0.06 | 0.05 | 0.04 | 0.03 | 0.02 | 0.02 | 0.02 | 0.01 | 0.01 | 0.01 | 0.01 |
| Cumulative Proportion | | 0.2 | 0.37 | 0.5 | 0.59 | 0.66 | 0.72 | 0.78 | 0.83 | 0.88 | 0.91 | 0.93 | 0.94 | 0.96 | 0.97 | 0.98 | 0.99 | 1 |

Loadings with absolute values greater than or equal to 0.25 are highlighted in bold.

**Table S5.** Scaling of the PCA factors for golden rocket frog calls on the discriminant functions, which show the relative importance of each PCA factor to the discriminant function, and variance explained by each discriminant function. All discriminant functions that cumulatively explained 100% of the variation are shown.

| | Discriminant function | | | | | | | | | | | |
|---|---|---|---|---|---|---|---|---|---|---|---|---|
| PCA factor | 1 | 2 | 3 | 4 | 5 | 6 | 7 | 8 | 9 | 10 | 11 | 12 |
| Factor 1 | -0.80 | 0.50 | 0.22 | **-0.49** | 0.03 | 0.01 | -0.13 | 0.06 | 0.00 | 0.01 | -0.09 | 0.10 |
| Factor 2 | **-1.71** | 0.16 | -0.25 | 0.21 | 0.02 | -0.03 | 0.09 | -0.03 | -0.02 | -0.05 | 0.05 | -0.15 |
| Factor 3 | -0.81 | **-1.24** | 0.44 | -0.01 | -0.05 | 0.09 | -0.05 | 0.03 | -0.04 | 0.03 | -0.08 | 0.08 |
| Factor 4 | -0.02 | -0.69 | **-0.97** | -0.48 | 0.07 | 0.08 | -0.21 | 0.05 | 0.08 | 0.00 | 0.17 | -0.06 |
| Factor 5 | 0.26 | -0.11 | -0.31 | -0.32 | 0.01 | -0.02 | 0.18 | -0.01 | -0.24 | -0.26 | 0.11 | 0.02 |
| Factor 6 | -0.08 | 0.40 | -0.38 | 0.22 | -0.60 | 0.44 | -0.20 | 0.06 | -0.22 | 0.24 | -0.07 | 0.02 |
| Factor 7 | 0.44 | 0.00 | 0.53 | -0.22 | -0.21 | 0.20 | -0.19 | 0.20 | 0.15 | -0.20 | -0.01 | **-0.63** |
| Factor 8 | 0.10 | -0.16 | -0.51 | -0.13 | -0.36 | 0.08 | 0.39 | 0.08 | 0.16 | -0.30 | -0.60 | 0.12 |
| Factor 9 | 0.11 | 0.03 | -0.33 | 0.35 | 0.14 | -0.36 | -0.75 | -0.09 | 0.12 | -0.04 | -0.28 | 0.10 |
| Factor 10 | -0.34 | 0.01 | -0.28 | 0.21 | 0.07 | 0.03 | 0.08 | 0.66 | 0.56 | 0.40 | -0.04 | 0.10 |
| Factor 11 | 0.06 | 0.08 | 0.00 | -0.33 | 0.37 | 0.25 | 0.10 | -0.34 | -0.50 | 0.36 | -0.40 | -0.07 |
| Factor 12 | 0.09 | -0.44 | -0.18 | -0.45 | -0.65 | -0.58 | 0.00 | -0.66 | -0.03 | 0.69 | -0.17 | -0.41 |
| Factor 13 | -0.12 | -0.13 | 0.18 | -0.17 | **-0.92** | **-1.40** | 0.04 | 0.61 | -0.27 | -0.13 | 0.19 | 0.08 |
| Factor 14 | -0.26 | -0.11 | 0.12 | -0.12 | -0.43 | -0.04 | 0.14 | -0.13 | -0.31 | 0.23 | 0.45 | 0.19 |
| Factor 15 | 0.10 | -0.30 | -0.42 | 0.39 | 0.52 | -0.44 | -0.23 | -0.08 | -0.44 | -0.22 | **-0.80** | -0.58 |
| Factor 16 | -0.85 | 0.16 | 0.38 | -0.10 | -0.85 | 0.09 | -0.12 | **-1.44** | **1.35** | -0.57 | 0.29 | 0.33 |
| Factor 17 | -0.40 | 0.09 | 0.11 | 0.76 | -0.58 | 0.50 | **-1.08** | 0.19 | -0.73 | **-1.10** | 0.30 | 0.53 |
| Singular value | 18.42 | 11.46 | 8.39 | 5.82 | 3.96 | 3.57 | 3.13 | 2.49 | 2.22 | 1.81 | 1.70 | 1.63 |
| Proportion of variance | 0.53 | 0.21 | 0.11 | 0.05 | 0.02 | 0.02 | 0.02 | 0.01 | 0.01 | 0.01 | 0.00 | 0.00 |
| Cumulative proportion | 0.53 | 0.74 | 0.85 | 0.90 | 0.93 | 0.95 | 0.96 | 0.97 | 0.98 | 0.99 | 0.99 | 1.00 |

The largest loading for each discriminant function is highlighted in bold.

**Table S6.** Loadings of measured Kai rocket frog call properties on principal components and the variance explained by each component.

| Call property | | PCA Factor | | | | | | | | | | |
|---|---|---|---|---|---|---|---|---|---|---|---|---|
| | | 1 | 2 | 3 | 4 | 5 | 6 | 7 | 8 | 9 | 10 | 11 |
| Call temporal properties | Call duration | **-0.26** | **0.31** | **0.36** | -0.05 | 0.08 | 0.00 | 0.09 | -0.11 | -0.06 | 0.03 | 0.10 |
| | Call interval | -0.07 | 0.16 | -0.15 | **0.60** | **0.29** | -0.02 | -0.08 | 0.02 | -0.03 | 0.00 | -0.05 |
| | Call period | -0.07 | 0.16 | -0.15 | **0.59** | **0.29** | -0.02 | -0.08 | 0.02 | -0.03 | 0.00 | -0.05 |
| Pulse duration | Pulse 1 | **-0.26** | **0.31** | -0.14 | -0.23 | 0.15 | **0.27** | 0.12 | -0.08 | 0.18 | -0.12 | **-0.58** |
| | Pulse 2 | **-0.26** | **0.31** | -0.19 | -0.11 | 0.03 | **0.38** | 0.04 | **-0.28** | -0.24 | 0.10 | **0.56** |
| Pulse interval | Pulse 1 | -0.08 | 0.12 | **0.57** | 0.10 | 0.01 | **-0.27** | 0.02 | 0.03 | -0.05 | 0.04 | 0.12 |
| Pulse period | Pulse 1 | -0.19 | **0.25** | **0.48** | -0.01 | 0.08 | -0.13 | 0.08 | -0.01 | 0.03 | -0.02 | -0.14 |
| Pulse rise time | Pulse 1 | **-0.38** | -0.18 | -0.11 | -0.12 | 0.15 | -0.16 | -0.13 | **-0.47** | 0.08 | 0.00 | **-0.26** |
| | Pulse 2 | **-0.35** | -0.21 | -0.10 | -0.05 | 0.23 | 0.07 | **0.46** | 0.17 | -0.10 | -0.04 | **0.25** |
| Pulse fall time | Pulse 1 | 0.21 | **0.39** | 0.01 | -0.04 | -0.04 | **0.34** | 0.22 | **0.46** | 0.04 | -0.07 | -0.13 |
| | Pulse 2 | 0.22 | **0.38** | 0.00 | -0.01 | -0.22 | 0.14 | **-0.44** | **-0.32** | -0.03 | 0.09 | 0.04 |
| Pulse 50% rise time | Pulse 1 | **-0.31** | 0.13 | -0.13 | -0.16 | -0.02 | -0.21 | **-0.48** | **0.37** | 0.06 | **-0.62** | 0.20 |
| | Pulse 2 | **-0.35** | 0.10 | -0.17 | -0.11 | -0.13 | -0.17 | -0.24 | **0.42** | -0.03 | **0.73** | -0.09 |
| Pulse 50% fall time | Pulse 1 | 0.07 | **0.32** | **-0.26** | 0.01 | -0.12 | **-0.43** | **0.33** | -0.13 | **0.66** | 0.05 | **0.25** |
| | Pulse 2 | 0.08 | **0.27** | **-0.27** | -0.06 | -0.19 | **-0.48** | **0.28** | -0.08 | **-0.66** | -0.13 | -0.19 |
| Dominant frequency | Pulse 1 | **0.26** | 0.08 | -0.03 | **-0.29** | **0.55** | -0.13 | -0.08 | 0.03 | -0.04 | 0.09 | 0.02 |
| | Pulse 2 | **0.27** | 0.08 | -0.02 | **-0.26** | **0.55** | -0.15 | -0.11 | 0.02 | -0.04 | 0.10 | 0.11 |
| Variance | | 4.25 | 3.36 | 2.56 | 1.97 | 1.61 | 1.00 | 0.81 | 0.56 | 0.35 | 0.26 | 0.22 |
| Proportion of Variance | | 0.25 | 0.20 | 0.15 | 0.12 | 0.09 | 0.06 | 0.05 | 0.03 | 0.02 | 0.02 | 0.01 |
| Cumulative Proportion | | 0.25 | 0.45 | 0.60 | 0.71 | 0.81 | 0.87 | 0.91 | 0.95 | 0.97 | 0.98 | 1.00 |

Loadings with absolute values greater than or equal to 0.25 are highlighted in bold.

**Table S7.** Scaling of the PCA factors for Kai rocket frog calls on the discriminant functions, which show the relative importance of each PCA factor to the discriminant function, and variance explained by each discriminant function. All discriminant functions that cumulatively explained 100% of the variation are shown.

| | Discriminant function | | | | | | | | |
|---|---|---|---|---|---|---|---|---|---|
| PCA factor | 1 | 2 | 3 | 4 | 5 | 6 | 7 | 8 | 9 |
| Factor 1 | **-1.13** | 0.29 | -0.14 | 0.23 | -0.17 | 0.26 | -0.02 | 0.03 | -0.04 |
| Factor 2 | 0.59 | -0.51 | -0.35 | 0.61 | 0.01 | 0.20 | 0.01 | -0.02 | 0.07 |
| Factor 3 | -0.49 | -0.25 | 0.67 | 0.42 | 0.52 | -0.06 | 0.04 | 0.03 | -0.10 |
| Factor 4 | 0.21 | 0.74 | 0.13 | 0.31 | -0.03 | -0.07 | 0.19 | -0.54 | 0.18 |
| Factor 5 | -1.01 | **-1.18** | -0.04 | -0.27 | -0.25 | -0.15 | 0.19 | -0.32 | 0.06 |
| Factor 6 | 0.80 | -0.21 | **1.02** | -0.05 | -0.73 | 0.58 | -0.28 | -0.03 | -0.10 |
| Factor 7 | 0.45 | 0.20 | -0.35 | -0.32 | 0.41 | 0.40 | 0.50 | 0.11 | -0.49 |
| Factor 8 | -0.36 | 0.13 | -0.39 | 0.28 | 0.13 | -0.50 | **-1.13** | 0.01 | 0.00 |
| Factor 9 | 0.15 | -0.15 | 0.10 | -0.49 | 0.36 | 0.21 | -0.42 | -0.42 | 0.28 |
| Factor 10 | 0.47 | -0.13 | -0.35 | 0.30 | -0.33 | -0.42 | -0.34 | -0.78 | **-1.63** |
| Factor 11 | 0.30 | 0.32 | 0.43 | **1.06** | **-1.51** | **-1.43** | 0.73 | 0.76 | -0.12 |
| Singular value | 13.44 | 8.81 | 6.39 | 5.94 | 4.95 | 3.61 | 2.79 | 1.74 | 1.45 |
| Proportion of variance | 0.47 | 0.20 | 0.11 | 0.09 | 0.06 | 0.03 | 0.02 | 0.01 | 0.01 |
| Cumulative proportion | 0.47 | 0.67 | 0.77 | 0.87 | 0.93 | 0.96 | 0.98 | 0.99 | 1.00 |

The largest loading for each discriminant function is highlighted in bold.

**Table S8.** Loadings of measures of aggression on the aggression indices derived from principal components transformations of aggressive responses for each species, as well as variance explained by each index.

| | Golden rocket frog aggression index (PC1) | Kai rocket frog aggression index (PC1) |
|---|---|---|
| Aggressive calls | 0.51 | 0.53 |
| Pseudo-aggressive calls | 0.08 | 0.53 |
| Approach distance | 0.63 | 0.52 |
| Closest position to speaker | 0.57 | 0.39 |
| Variance | 1.81 | 2.08 |
| Proportion of variance | 0.43 | 0.52 |
